# Direct detection of RNA repair by nanopore sequencing

**DOI:** 10.1101/2022.05.29.493267

**Authors:** Laura K. White, Saylor M. Strugar, Andrea MacFadden, Jay R. Hesselberth

## Abstract

Ligation by plant and fungal RNA ligases yields an internal 2′-phosphate group on each RNA ligation product. In budding yeast, this covalent mark occurs at the splice junction of two targets of ligation: intron-containing tRNAs and the messenger RNA *HAC1*. The repertoire of RNA molecules repaired by RNA ligation has not been explored due to a lack of unbiased approaches for identifying RNA ligation products. Here, we define several unique signals produced by 2′-phosphorylated RNAs during nanopore sequencing. A 2′-phosphate at the splice junction of *HAC1* mRNA inhibits 5′→3′ degradation, enabling detection of decay intermediates in yeast RNA repair mutants by nanopore sequencing. During direct RNA sequencing, intact 2′-phosphorylated RNAs produce diagnostic changes in nanopore current properties and base calling features, including stalls produced as the modified RNA translocates through the nanopore motor protein. These approaches enable directed studies to identify novel RNA repair events from multiple species.

## Introduction

RNA ligases play a key role in the repair of endonuclease-damaged RNA ^1^. In fungi and plants, Trl1 joins the exons of intron-containing tRNAs after intron excision by the tRNA splicing endonuclease ^2^. The prokaryotic and metazoan RNA ligase RtcB catalyzes repair of endonuclease-cleaved tRNAs to effect tRNA splicing ^3^, and like Trl1, activates the unfolded protein response in animals ^4,5^. While RtcB ligation yields a 3′-5′ phosphodiester linkage ^6^, plant and fungal ligases produce a 2′-phosphate at the ligation junction, marking the site of RNA repair ^7^.

The 2′-phosphate products of Trl1 ligation are subsequently removed by the 2′-phosphotransferase Tpt1 ^8^. To investigate the role of RNA repair in eukaryotes, we previously generated a genetic bypass of *S. cerevisiae TRL1* and *TPT1* by expressing intronless versions of all ten intron-containing yeast tRNAs ^9^, and used these cells to investigate how RNA repair enzymes regulate the non canonical splicing of *HAC1*, a messenger RNA encoding a transcription factor that activates the unfolded protein response (UPR) ^4^. We showed that the 2′-phosphate on ligated *HAC1* mRNA in *xrn1*Δ *tpt1*Δ cells protects its 3′-exon from degradation by Xrn1, raising the possibility that the 2′-phosphates produced by Trl1 ligation might directly inhibit this exonuclease ^10^.

Nanopore sequencing enables simultaneous analysis of the sequence and modifications of native RNA molecules. In budding yeast, this approach has been used to identify sites of methylation (m^6^A and 2′-O-methylation) and pseudouridylation in mRNAs, rRNA, and some non-coding RNAs ^11–14^. Technical hurdles associated with direct RNA sequencing of tRNA, the most highly modified class of RNA, were recently overcome by a strategy used to sequence *E. coli* tRNA ^15^. We applied these and other approaches to develop a strategy to identify 2′-phosphorylated RNAs produced during RNA repair.

## Results

### 2′-phosphate modifications are sufficient to inhibit 5′→3′ exoribonucleases

Our studies of *HAC1* mRNA processing in the UPR revealed that 2′-phosphorylated RNA processing intermediates are resistant to 5′→3′ exonucleolytic degradation by Xrn1^10^ (**Fig. 1a**), resonating with the previous finding that 2′-phosphates inhibit a bacterial 3′→5′ exonuclease ^16^. This effect is generalizable to the other known substrates of Trl1, intron-containing tRNAs (**Fig. 1b**). We reconstituted 2′-phosphate-mediated exonuclease stalling *in vitro* by digesting synthetic RNAs with 2′-phosphate ^17^ or 2′-O-methyl modifications with recombinant 5′→3′ exonucleases ^18,19^ (**Fig. 1c, d**), confirming that 2′-phosphates are sufficient to inhibit multiple 5′→3′ exonucleases, and provide more robust protection than a 2′-O-methyl group. DxoI and RNase J1 stall 1-3 nt upstream of the 2′-phosphorylated site, while Xrn1 is capable of digesting an RNA up to the 2′-phosphorylated nucleotide. We found that these 5′,2′-phosphorylated RNAs are poor substrates for ligation (**Fig. S1**), precluding a standard approach to identify 5′-ends ^20^.

**FIGURE 1.**
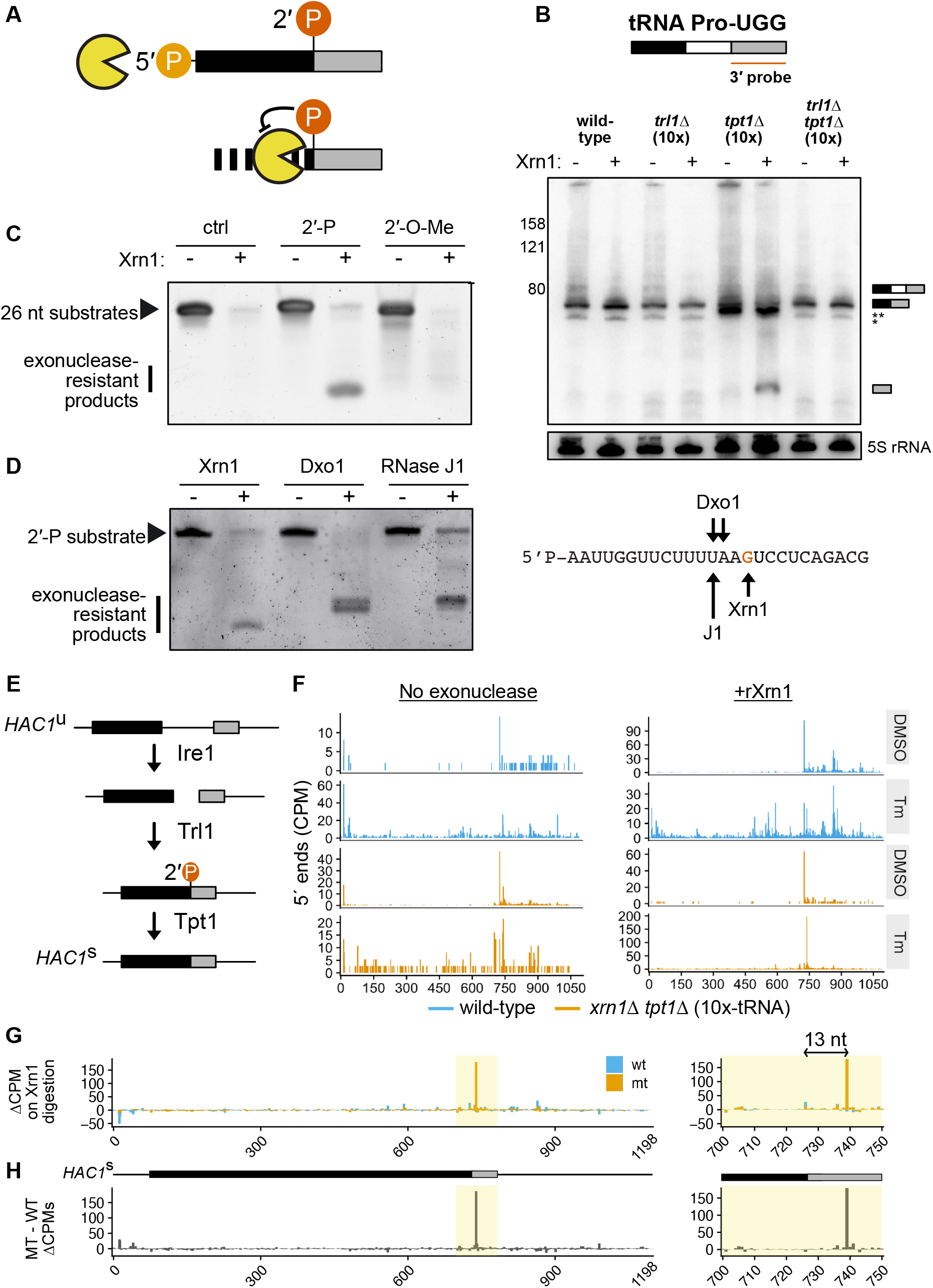
*In vitro* digestion with recombinant exonucleases reveals sites of 2′-phosphorylation. (**A**) Digestion of 2′-phosphorylated RNA by Xrn1 yields a product that inhibits further degradation. (**B**) The 2′-phosphate on tRNA-Pro-UGG can be detected by northern blotting for 3′-exon in total RNA from RNA repair strains treated with and without recombinant Xrn1 *in vitro*. In the lower panel, the same membrane has been stripped and reprobed for 5S rRNA as a loading control. Asterisks correspond to previously described but uncharacterized Pro-TGG splicing intermediates ^9,37^. (**C**) Unmodified, 2′-phosphorylated, and 2′-O-methylated RNA substrates show zero, substantial, and trace resistance to Xrn1 digestion, respectively. (**D**) A 2′-phosphate inhibits decay by the 5′→3′ exonucleases Xrn1, Dxo1, and RNase J1. The location of the 2′-phosphorylated linkage is indicated by the red “G” in the substrate at right, with inferred stalling sites for each exonuclease marked with arrows. (**E**) In budding yeast, the unfolded protein response (UPR) initiates a noncanonical splicing event on the mRNA *HAC1* during which the endonuclease Ire1 excises the *HAC1* intron (thin line), followed by exon ligation by Trl1 (black and gray boxes), generating an internal 2′-phosphate at the splice junction that is subsequently removed by the 2′-phosphotransferase Tpt1. (**F**) RNA from WT and *xrn1Δ tpt1Δ (10x-tRNA)* cells was treated with DMSO or tunicamycin and left untreated (left panel) or enzymatically and digested (right panel) followed by nanopore sequencing. 5′-end alignments of reads are expressed in counts per million reads (CPM). (**G**) Subtraction of undigested background signal from signal from the Tm-treated, rXrn1-digested libraries in (**F**), yields a predominant 5′-end signal on *HAC1*^s^ at the splice junction (yellow box). At right, an enhanced view of this region shows the major peak in wild-type cells is located precisely at the exon-exon junction (position 727), whereas the end signal in the mutant is 13 nucleotides downstream, an offset consistent with premature termination of base calling when a *bona fide* 5′ RNA end exits a nanopore. (**H**) The wild-type rXrn1 enrichment scores above have been subtracted from the mutant scores at each nucleotide position.

### Nanopore sequencing of Xrn1-degraded RNA identifies 2′-phosphates produced during *HAC1* splicing

We asked whether Xrn1 treatment of total RNA could enhance the 5′-end signal on Trl1 substrates in RNA sequencing libraries. We treated wild-type and *xrn1*Δ *tpt1*Δ cells with either DMSO or tunicamycin, which inhibits N-linked glycosylation and induces the UPR, triggering the endonuclease Ire1 to excise the intron from unspliced, cytoplasmic *HAC1* pre-mRNA (*HAC1*^u^) (**Fig. 1e**). The exon halves are ligated by Trl1 ^4^, yielding a spliced mRNA molecule (*HAC1*^s^) bearing an internal 2′-phosphate at the ligation junction that is removed by the 2′-phosphotransferase Tpt1 ^8^. In *tpt1Δ* cells, the 2′-phosphate stabilizes *HAC1*^s^ against 5′→3′ exonucleolytic degradation ^10^.

We treated total RNA with mRNA decapping enzyme ^21^ to expose 5′-monophosphate ends and digested RNAs with recombinant Xrn1 (rXrn1) and then prepared libraries for direct mRNA nanopore sequencing (**Table S1**). After base calling, 5′-read ends from each library were aligned to a budding yeast transcriptome containing both *HAC1*^u^ and *HAC1*^s^. Consistent with previous results ^10^, *HAC1* 3′-exon termini accumulate in *xrn1*Δ *tpt1*Δ cells even in the absence of tunicamycin, and are enriched upon Xrn1 treatment (**Fig. 1f**). Comparative analysis of signal from tunicamycin-treated samples confirmed this enrichment is strongly dependent on genotype; the largest change in 5′-end coverage occurs in the *xrn1Δ tpt1Δ* mutant and the peak is located 13 nt downstream of the *HAC1* splice junction (**Fig. 1g**), an offset consistent with premature termination of base calling when a *bona fide* 5′-end exits the pore ^22^. Subtracting the wild-type control from the mutant signal (**Fig. 1h**) showed that the largest genotype-specific change in 5′-end signal upon exonuclease treatment is the peak at nucleotide 740, consistent with 2′-phosphate mediated inhibition of Xrn1 degradation with single nucleotide resolution.

### RNA repair generates localized base calling errors on *HAC1*

We next asked whether we could directly detect 2′-phosphates produced during *HAC1* splicing without the use of exonuclease treatment. In principle, direct RNA sequencing can be used to detect any RNA modification that produces an alteration in current as the modified nucleotide passes through the nanopore sensor ^23^. These distortions affect the accuracy of base calling, generating base calling “errors” in the form of mismatches at or near the modified position. However, detection and discrimination of modified nucleotides can be strongly affected by (i) the RNA modification type, (ii) the sequence context in which the modification occurs, and (iii) the stoichiometry of the modification ^11,13^.

Inspection of all reads aligning to *HAC1*^s^ from tunicamycin-treated cells identified several positions with high rates of mismatched nucleotides, suggesting the presence of an RNA modification (**Fig. 2a**). While several of these exceed rates of 20% mismatching (e.g., 20% of reads spanning a site contain a mismatch at that position) in both the wild-type and mutant sample, a unique pattern of base calling errors occurs in the *xrn1*Δ *tpt1*Δ mutant, and is centered on the *HAC1*^s^ splice junction. At this location, the 2′-phosphorylated guanosine is only mis-called 3% of the time, but mismatch rates for the four flanking nucleotides in the 5mer centered on this position range from 14-29% (**Fig. 2b and Fig. S2**). These base calling errors were accompanied by a localized drop in the per-nucleotide quality score at the *HAC1*^s^ ligation junction (**Fig. 2c**), with the lowest values at the 2′-phosphorylated guanosine (position 727) in *xrn1*Δ *tpt1*Δ cells.

**FIGURE 2.**
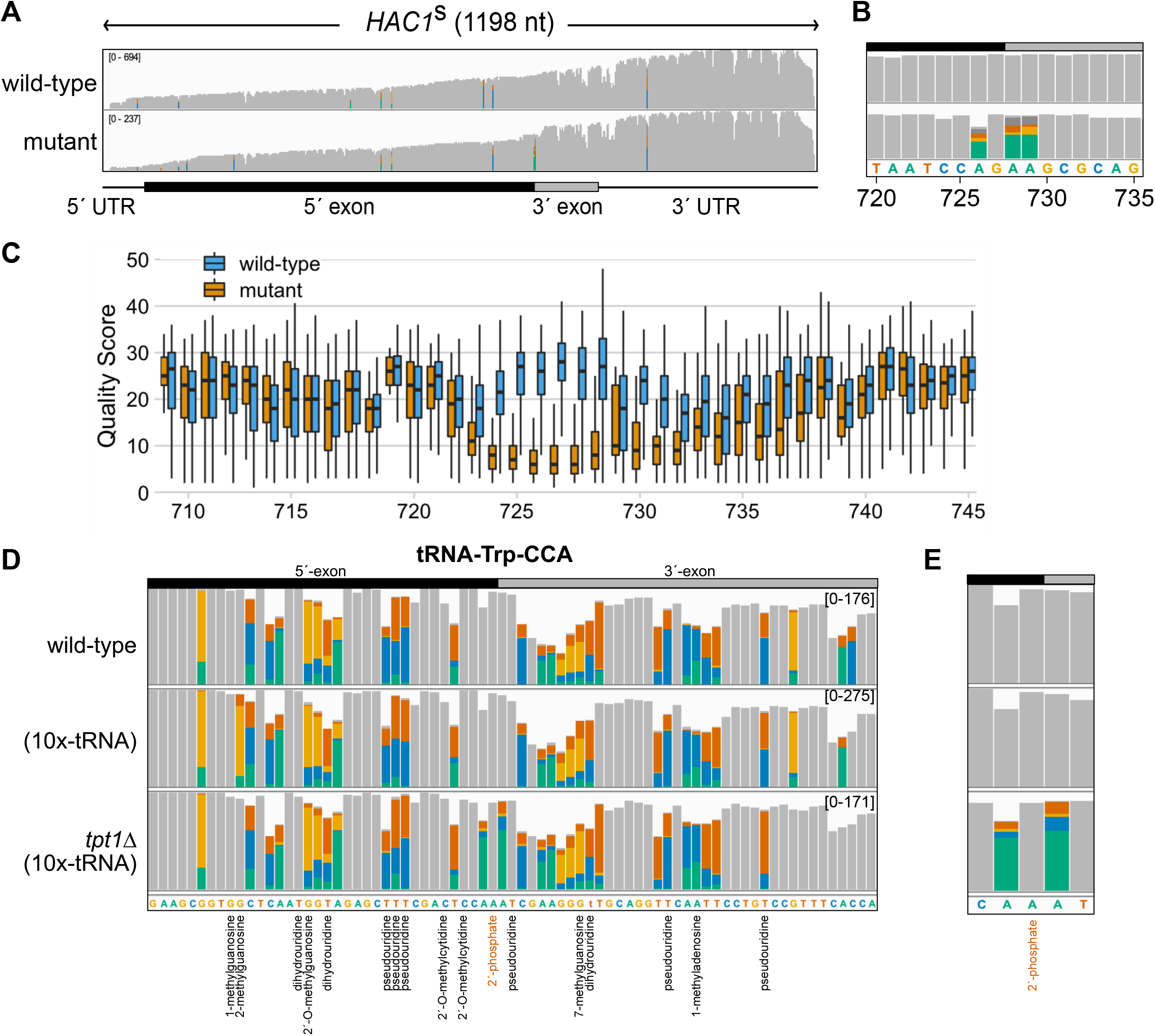
Nanopore sequencing enables direct detection of known 2′-phosphates deposited by Trl1 ligation *in vivo*. (**A**) Coverage of reads from Tm-treated, undigested mRNA libraries from wild-type and *xrn1Δ tpt1Δ* (10x-tRNA) cells (“mutant”) aligned to *HAC1*^s^. Bar heights quantify read coverage (y-limits, upper left). Colored bars represent positions with >20% mismatching to the reference base; gray bars indicate positions that did not exceed this threshold. (**B**) An enhanced view of the splice junction at right shows mutant-specific mismatches at nucleotides flanking the 2′-phosphorylated guanosine. (**C**) Base calling quality scores over this region show a reduction in base calling in mutant RNA centered on the 2′-phosphorylated position. (**D**) The 2′-phosphate on tRNA-Trp-CCA can be detected by anticodon-proximal mismatches in tRNA sequencing reads mapping to the mature tRNA reference. Each colored bar represents a position with >20% mismatching to the reference base; gray bars indicate positions which did not exceed this threshold. Modified positions reported in MODOMICS are annotated below the reference sequence, with the 2′-phosphate indicated in red text. Values in brackets represent the range of coverage in raw read counts across the entire tRNA. (**E**) Close-up of the 5 nt centered on the 2′-phosphorylated adenosine located at the tRNA-Trp-CCA splice junction.

### Complex modifications of tRNA complicate detection of RNA repair signals

We wondered whether such signals were present on intron-containing tRNAs ligated by Trl1. Budding yeast tRNAs contain an average of 13 modifications per tRNA species deposited across 36 different modified positions ^24^. The nucleotide immediately 3′ of the tRNA anticodon, termed the “hypermodified base”, is modified >55% of the time in budding yeast ^25^; on intron-containing tRNAs, this same nucleotide is also 2′-phosphorylated during tRNA splicing (**Fig. S3a**). Our examination of the modifications present within all ten intron-containing budding yeast tRNAs in the 5 nt window surrounding this position found that only tryptophan tRNAs lack an annotated modification at the hypermodified base (**Fig. S3b,c**); therefore, we focused on Trp-CCA to determine whether it was possible to directly detect 2′-phosphates on tRNAs. We adapted a method^15^ to ligate adapters to purified tRNA in preparation for direct RNA sequencing. While the average spliced tRNA in these libraries contained >28 positions where mismatch rates exceeded 20% (**Fig. S3d**), we observed a clear *TPT1*-dependent base calling error signature on Trp-CCA tRNA (**Fig. 2d,e**), consisting of a genotype-specific increase in mismatching at nucleotides flanking the 2′-phosphorylated position.

### Covalent marks of RNA repair produce distortions in nanopore signal intensity

While base calling errors are commonly used to detect modifications from direct RNA-seq data, a more direct method is to examine RNA modifications on raw nanopore signal intensity. Both of these approaches have been exploited in the development of computational tools to detect and quantify RNA modifications ^26^. We used Nanopolish ^27^ to annotate the reads aligning to *HAC1*^s^ with the level of ionic current observed as individual ribonucleotides pass through the center of the nanopore sensor. The largest difference in mean current intensity between our wild-type and mutant samples was located precisely at the 2′-phosphorylated guanosine (**Fig. 3a**). Plotting per-read signal revealed substantial divergence in current intensity within a 15 nucleotide window surrounding the *HAC1*^s^ splice junction (**Fig. 3b**). Principal components analysis of the per-read current intensities over this region and a density plot of all current intensities within the same window show significant differences between the wild-type and mutant signals (**Fig. 3c,d**) indicating that deviation in current intensity provides useful information to detect 2′-phosphate modifications.

**FIGURE 3.**
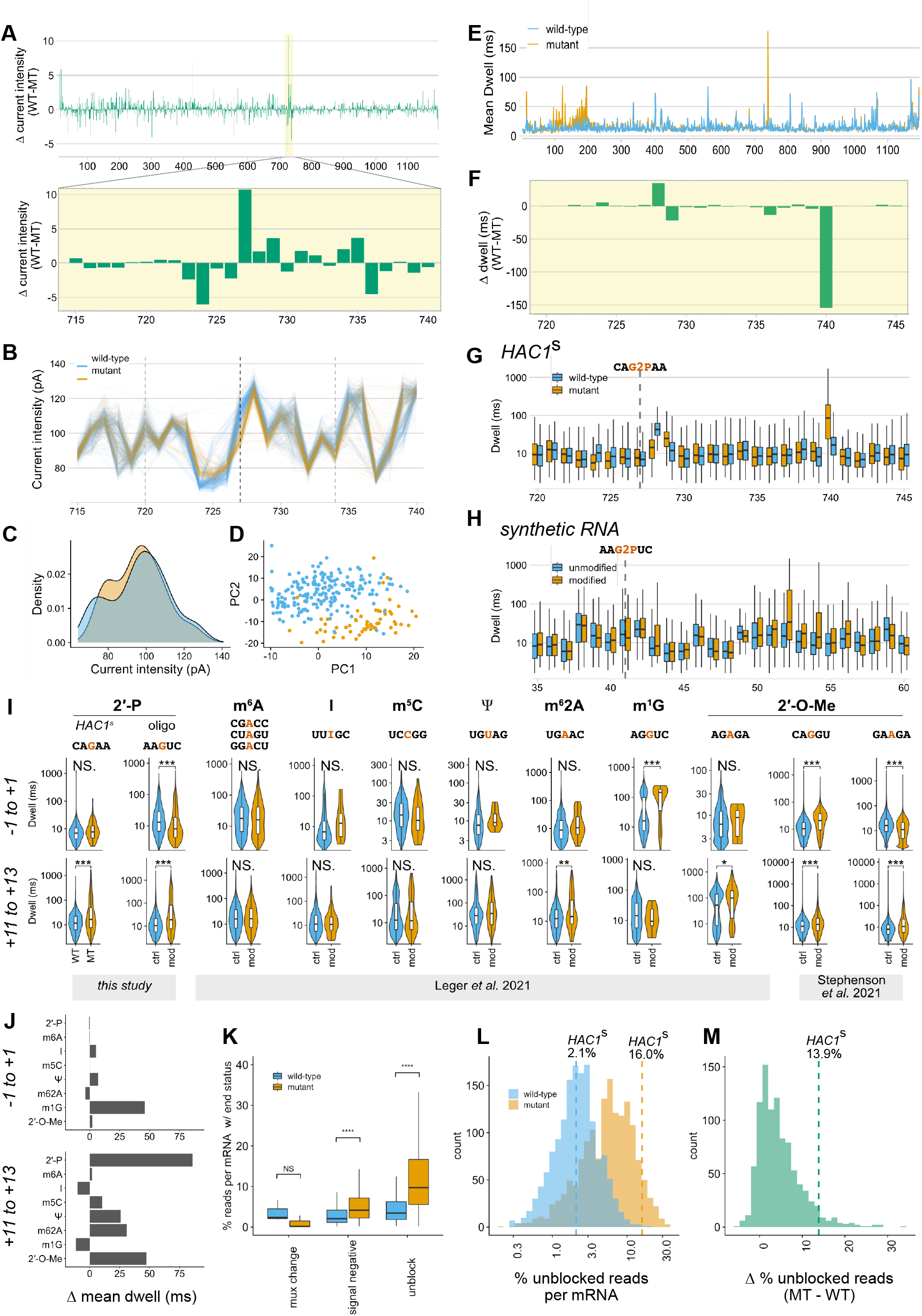
Current and base calling features of 2′-phosphorylation and other RNA modifications. (**A**) Change in mean current intensity across the spliced *HAC1*^*s*^ transcript (upper panel) and at the splice junction (lower panel). (**B**) Per-read current intensities at the *HAC1*^*s*^ splice junction. The dotted gray lines define a 15 nt window corresponding to the approximate footprint of the nanopore and docked motor protein when the 2′-phosphorylated nucleotide (vertical dashed black line, position 727) is centered in the reader head of the nanopore. (**C**) Distribution of current intensities across all reads within this 15 nt window. (**D**) Principal components analysis of current intensity values across the 15 nt window surrounding the splice site. (**E**) Mean dwell time in milliseconds for each nucleotide across *HAC1*^s^. (**F**) Difference in dwell time between WT and mutant samples within the region surrounding the *HAC1* splice junction. (**G**) Distribution of per-read dwell times in WT and mutant samples over this same region, with the *HAC1* splice junction indicated by a vertical dashed line. (**H**) Difference in dwell time on reads aligning to a synthetic RNA within the region surrounding the 2′-phosphorylated or unmodified position (vertical dashed line). (**I**) Violin plots of the distribution of per-read dwell times within a 3 nt window surrounding a modified position (upper row) or a 3 nt window +11 nt in the 3′ direction on the same RNA substrates. Overlaid box plots define the median and interquartile range. Above each stacked pair of plots, the sequence context(s) for each modification are indicated with the modified nucleotide in red. (**J**) The difference in dwell times for all modifications in (**I**) was calculated by subtracting the mean dwell for modified RNA vs. the mean dwell from the unmodified libraries, over the same 3 nt windows. 2′-phosphate, m^6^A and 2′-O-methyl means were calculated across multiple sequence contexts. (**K**) Box plots showing the percent of aligned reads per mRNA in UPR-stressed cells that did not report a successful positive signal at termination, with the alternative end reasons on the x axis. (**L**) Histogram displaying the percent of reads per mRNA in WT or mutant libraries with the “unblock” end status. In this plot, only mRNAs with ≥30 total reads are visualized. Vertical dashed blue and orange lines indicate the percent of unblocked reads aligning to *HAC1*^*s*^ in the WT and mutant samples, respectively. (**M**) For each mRNA in (**L**), the percent of unblocked reads in the WT sample was subtracted from the corresponding percentage in the mutant sample. The location of *HAC1*^*s*^ on this histogram is indicated with a vertical dashed green line.

### RNA repair generates detectable stalls during pore translocation

An additional direct output of nanopore sequencing is dwell, the residence time of an individual nucleotide within the nanopore sensor. Previous reports described varying utility of dwell time for detecting specific RNA modifications ^11,13,14,28^. In *xrn1*Δ *tpt1*Δ cells, we observed a >150 millisecond increase in dwell 13 nt 3′ of the *HAC1*^s^ ligation junction (**Fig. 3e,f**). As RNA is threaded into the nanopore in the 3′→5′ direction by a helicase, we infer that this stalling occurs before the 2′-phosphate has entered the nanopore channel and reflects interactions between the modification and the motor protein, similar to 2′-O-methyl and pseudouridine modifications ^14,28^. To validate this finding, we examined the distribution of dwell times in RNA sequencing reads over the *HAC1* splice junction (**Fig. 3g**), and performed direct RNA-seq on a synthetic RNA substrate, which also showed a 128 millisecond increase in mean dwell at position +11 nt (**Fig. 3h and S4**).

To contextualize the magnitude and specificity of this effect, we re-analyzed nanopore dwell times on modified and unmodified oligonucleotides ^13^ and at select modified sites on native budding yeast rRNA vs. an *in vitro* transcribed control ^14^. Among eight different RNA modifications, only 2′-phosphorylation, N6,N6-dimethylation (m^6^2A), and 2′-O-methylation produced statistically significant changes in dwell times at 3′ distal sites located +11-13 nt from the RNA modification (**Fig. 3i**). 2′-phosphates generated the largest increases in mean dwell time of all modifications analyzed (**Fig. 3j**), suggesting that dwell time is a robust signal for the detection of RNA repair events. While this analysis covers only a small repertoire of sequence contexts, our results suggest that approaches for *de novo* detection of RNA modifications in direct RNA-seq may be improved by incorporation of distal stalling signals produced as RNA modifications traverse the nanopore motor protein (**Fig. S5**).

Nanopore sequencing devices can manipulate individual nanopores wherein nucleic acids become stalled by reversing the local electric potential in an attempt to eject the stalled molecule. **Fig. 3k** shows the percentage of aligned reads per mRNA that terminated unexpectedly. We found a statistically significant increase in reads terminated due to negative signal and unblocking events in *xrn1*Δ *tpt1*Δ cells treated with tunicamycin, suggesting additional products of RNA repair. To explore this further, we selected mRNAs with sufficient coverage, and calculated the percent of reads per mRNA with this “unblock” status. We found a 14% increase in reads aligning to *HAC1*^s^ that terminated in an unblocking event in *xrn1Δ tpt1Δ* cells compared to wild-type (**Fig. 3l**). We calculated the strain-specific difference in percent of reads terminated by unblocking for all remaining mRNAs, and found 25 mRNAs with higher numbers of unblocked reads than *HAC1*^s^ (**Fig. 3m**). However, further inspection of these mRNAs did not identify signals consistent with site specific 2′-phosphorylation, indicating dwell alone is not sufficient to identify repair events.

In principle, 170 RNA modifications ^25^ are detectable by direct RNA sequencing. However, the specific signals produced by a modification in all RNA sequence contexts and the threshold of detection for a given modification require synthetic and genetic controls, as well as high depth sequencing coverage ^13^. Even high stoichiometry modifications (e.g., rRNA 2′-O-methylation) produce strong current intensity distortions in some sequence contexts but not in others ^11^. Accordingly, the development of tools capable of integrating and comparing multiple types of direct RNA-seq signals is an area of active investigation ^26^. In this work, we define a collection of signals produced by 2′-phosphates in nanopore sequencing that represent a diagnostic signature of RNA repair on the mRNA *HAC1*, and report the first direct RNA sequencing of eukaryotic tRNA. The recent finding that 2′-phosphates can also be deposited via phosphorylation ^29^ and an unknown role for animal 2′-phosphotransferase ^30^ highlight the potential for characterizing this RNA modification in additional biological contexts.

## Materials and Methods

### Exonuclease digestion of synthetic RNA substrates

All exonucleases were purified as described in ^19^. Equimolar amounts of the 26 nucleotide synthetic oligo substrates (Chemgenes) in **Table S2** were resuspended in 20 µL reactions containing either 1 µL of water or 1 µL of recombinant exonuclease at the following concentrations: RNaseJ1 = 1.1 mg/mL; Xrn1 = 0.8 mg/mL; Dxo1 = 0.68 mg/mL. These digestion reactions were incubated at 37 ºC for two hours in a standard buffer containing 100 mM NaCl, 50 mM Tris, 2 mM MgCl_2_ pH 7.9, and 1 mM DTT. An equal volume of 2X formamide loading dye was added to quench the reactions before running on a precast 15% TBE-urea acrylamide gel for 20 minutes at 125V, followed by 25 minutes at 175V. Gels were stained with SybrGold (ThermoFisher) and visualized on a BioRad Gel Doc imager.

### Yeast cell culture and RNA isolation

All cultures were inoculated from single colonies in synthetic drop-out media supplemented with relevant amino acids to maintain plasmid selection. Cells were grown at 30ºC to mid-log phase and, where indicated, treated for two hours with tunicamycin (Sigma-Aldrich, final concentration of 2.5 µg/mL) or an equal concentration of DMSO. After centrifugation, total RNA was isolated by hot acid phenol extraction. All yeast strains used in this study are listed in **Table S3**.

### Exonuclease-digested tRNA northern blots

Total RNA (15 µg) was digested in 100 mM NaCl, 50 mM Tris, 2 mM MgCl_2_ pH 7.9, and 1 mM DTT in 11 µL reactions containing 1 µL of rXrn1 at 1.1 mg/mL or 1 µL of DEPC H20 for 18 hours at 37ºC, and reactions stopped by addition of an equal volume of formamide loading dye. 5 µg of this total RNA input was loaded onto a precast 10% TBU acrylamide urea gel (Novex / ThermoFisher) and electrophoresed at 120 V for 15 minutes, followed by 170 V for 60 minutes. After staining and imaging, the gel was transferred at 3 mA/cm^2^ for 35 minutes to a charged nylon membrane (Hybond N+, GE) and UV crosslinked with a 120 mJ dose at 254 nm. The membrane was then blocked in Ambion ULTRAhyb-Oligo Buffer before incubating for 18 hours with a 5′-32P-labeled probe (**Table S2**) designed to hybridize to the 3′-exon of intron-containing tRNAs. After four 15 minute washes with 2X SSC/0.5% SDS washing buffer, the membrane was wrapped in plastic, exposed on a phosphor-imager screen, and subsequently imaged on a Typhoon 9400 (GE Healthcare). To re-probe, the membrane was washed twice in 2% SDS at 80ºC for 30 minutes per wash, re-blocked, and then incubated with a new probe as before.

### Preparation of exonuclease-degraded RNA for mRNA-seq

To prepare exonuclease-degraded mRNA, 200 µg of total RNA was decapped with mRNA decapping enzyme (New England Biolabs) for 1 hour at 37 ºC, ethanol precipitated, resuspended, and split into two 20 µL reactions in the buffer described above, with or without 2 µL of recombinant Xrn1 (1.1 mg/mL). Decapped RNA was incubated at 37 ºC for 5-18 hours, and degraded samples with RIN scores ranging from 1.5 - 4.5 as measured by Tapestation using an Agilent High Sensitivity ScreenTape were selected for further library preparation. The resulting samples were treated with phenol chloroform and ethanol precipitated before poly(A) selection using Dynabeads Oligo(dT)25 mRNA isolation beads (ThermoFisher Scientific). Yields were assayed by Nanodrop, and 500 ng of the resulting poly(A) selected RNA was used as input for direct RNAseq (Oxford Nanopore, SQK-RNA002).

### Direct mRNA-seq Nanopore libraries

The remaining mRNA-sequencing libraries were prepared as above, without a decapping or exonuclease degradation step. Select mRNA-seq libraries were ligated to the four custom barcoded DNA adapters described in ^31^ in lieu of the commercial RTA adapter from Oxford Nanopore. Libraries were pooled after reverse transcription and subsequent Ampure XP bead cleanup to prevent cross-ligation of barcodes. Following sequencing and base calling, all reads were assigned to one of the four barcodes or an unknown barcode bin using the DeepLexiCon software, and fastq files were separated by barcode. Reads with a confidence score of 95% or higher probability of correct barcode assignment were used for downstream analysis.

### Synthetic RNA libraries

We used a splinted ligation strategy to generate 2′-phosphate-containing synthetic RNA substrates for nanopore sequencing. We designed a DNA splint to ligate the 2′-phosphorylated or unmodified 26 nucleotide synthetic oligo substrates to the 5′ oligo AB (to prevent 5′-end signal artifacts near the modified position) and to the 3′ oligo CD, which contains a polyadenylated end competent for ligation by the ONT RTA adapter. Ligation with T4 RNL2 (New England Biolabs) at a 3-fold molar excess of RNA substrates to DNA splint, DNase I treated, and then run on a 10% denaturing gel to purify the 86 nt product. Purified RNAs (300-400 ng) were used as input for direct RNA sequencing. All oligos aside from the 26 mers described previously were purchased from Integrated DNA Technologies, and are detailed in **Table S2**.

### tRNA sequencing

*S. cerevisiae* tRNAs were size-selected by gel purification from 100 µg of total RNA. The 60-100 nucleotide fraction was excised from an 8% TBU acrylamide gel, extruded through a hole punctured in a 0.5 ml centrifuge tube, and eluted overnight at 4 ºC with rotation in a buffer of 300 mM sodium acetate, 1 mM EDTA, and 0.1% SDS, followed by filtration through a 0.22 µM pore Costar Spin-X centrifuge tube filter and ethanol precipitation. Pellets were resuspended in 10 µL of DEPC H_2_0 and quantified by Nanodrop before ligating 1 µg of tRNA to splint adapters as previously described ^15^. Ligation products (120-180 nt) were gel-purified, resuspended in 23 µL of DEPC H_2_0 and ligated to the Nanopore RMX adapter. The remaining steps of the library prep were carried out as described in the Oxford Nanopore direct RNA sequencing protocol. To increase yield, half of each prepared library was loaded directly onto the MinION flow cell, and the remaining half stored at 4 ºC. After the number of actively sequencing pores on the flow cell dropped to less than 50% of the initial run state, the run was paused and the second half of the library loaded.

### Sequencing run conditions and base calling

Direct RNA-seq libraries were loaded onto R9.4.1 flow cells in a MinION sequencer connected to a laptop running MinKNOW software version 21.10.4. Runs were performed with live base calling turned on to monitor read quality and translocation speed in real time. All libraries were subsequently base called with Guppy version 5.0.16 with default CPU parameters using the high accuracy model (rna_r9.4.1_70bps_hac.cfg).

### Alignment references and mapping parameters

Poly(A) selected mRNA libraries were mapped to an *S. cerevisiae* transcriptome reference ^32^, which contains both pre- and post-spliced mRNA references (https://github.com/Jacobson-Lab/yeast_transcriptome_v5). Reads were aligned using Minimap2^33^ in a splice-aware fashion with specific parameters (minimap2 -ax splice -uf -k14). tRNA sequencing libraries were mapped to a custom *S. cerevisiae* tRNA reference containing a consensus sequence for each cytoplasmic tRNA species in budding yeast, available at https://github.com/hesselberthlab/RNARePore. Consensus sequences are largely derived from the first predicted tRNA in each anticodon family as listed on gtRNAdb (e.g., tRNA-Phe-GAA-1-1)^34^. All tRNAs in this reference have had CCA added to their 3′ ends, tRNA splint adapter sequences appended and prepended, and (where applicable) introns removed *in silico*. tRNA alignments and alignments to the splint ligated synthetic RNA reference were performed using BWA-MEM version 0.7.16a ^35^ with the parameters bwa mem -W 13 -k 6 -x ont2d.

### Analysis of base calling differences and quality scores

Aligned reads and per-nucleotide mismatches were visualized in IGV version 2.11.2. tRNA reads were visualized in IGV and annotated with all modifications present for each *S. cerevisiae* tRNA species in MODOMICS^25^. We used EpiNano-RMS (https://github.com/novoalab/nanoRMS/tree/master/epinano_RMS)^11^ to extract the frequency of mismatch, insertion, and deletion rates per position, as well as per nucleotide base calling quality scores.

### Analysis and visualization of raw Nanopore signals

We used the Nanopolish^27^ eventalign module to reannotate our aligned sequencing reads with raw signal information from FAST5 format. While Nanopolish has been previously described as less effective at annotating signals from sequencing reads containing modified RNA^11^, a commonly used alternative software tool, Tombo, is restricted to Minimap2 alignments^36^, making it incompatible with short read sequencing from our synthetic oligonucleotide standards and tRNA libraries. We therefore extracted positional current intensity and dwell time information with Nanopolish on a per-read basis. As the resulting Nanopolish outputs can be quite large, these files were pre-processed with custom bash scripts to select specific regions of interest for further analysis in R.

We used the same approach to re-analyze published sequencing data from modified RNA oligonucleotide standards^13^ and ribosomal RNA ^14^. For this analysis, we realigned each publicly available fastq dataset to its published reference sequence, and then annotated the aligned reads with their dwell times from the corresponding FAST5 files using Nanopolish. We used this information to interrogate the distribution of dwell times over a 3 nt window centered at position 0 (the modified nucleotide) and a second 3 nt window centered at position 12, and performed two-sided Kolmogorov–Smirnov testing to determine whether there was a statistically significant difference in the distribution of dwell times between each pair of samples.

In the process, we observed an important nuance in the default behavior of Tombo and Nanopolish when annotating positional current and dwell information. While Nanopolish outputs signals in kmer-space, assigning them to the 5′ end of the 5 nucleotide window centered on the base called nucleotide, Tombo outputs the same signals in nucleotide-space, with both tools beginning their count from position zero. In light of this, we added three nucleotides to the position of all signals output by Nanopolish. This Tombo behavior explains the discrepancy between the original reported dwell signal at +10 position in ^14^ and our reanalysis, which places the peak dwell on the same 2′-O-methylated rRNA sites at +11 nucleotides.

### Analysis of read termination status

Sequencing reads that do not terminate with the expected increase in observed current caused by a nucleic acid strand exiting the pore are assigned one of three alternative end reasons by the ONT MinKNOW software, depending on whether they were terminated during a routine scan of the flow cell (“mux change”), terminated due to an unexpected drop in observed current (“signal negative”), or terminated after a MinKNOW-initiated attempt to remove the obstruction (“unblock mux change”). We used the MinKNOW-produced sequencing_summary.txt files from tunicamycin-treated cells to annotate sequencing reads aligned with Minimap2 with the reason that each read terminated, and calculated the proportion of reads per mRNA assigned to each of these end statuses.

## Figure Legends

**SUPPLEMENTAL FIGURE 1.**
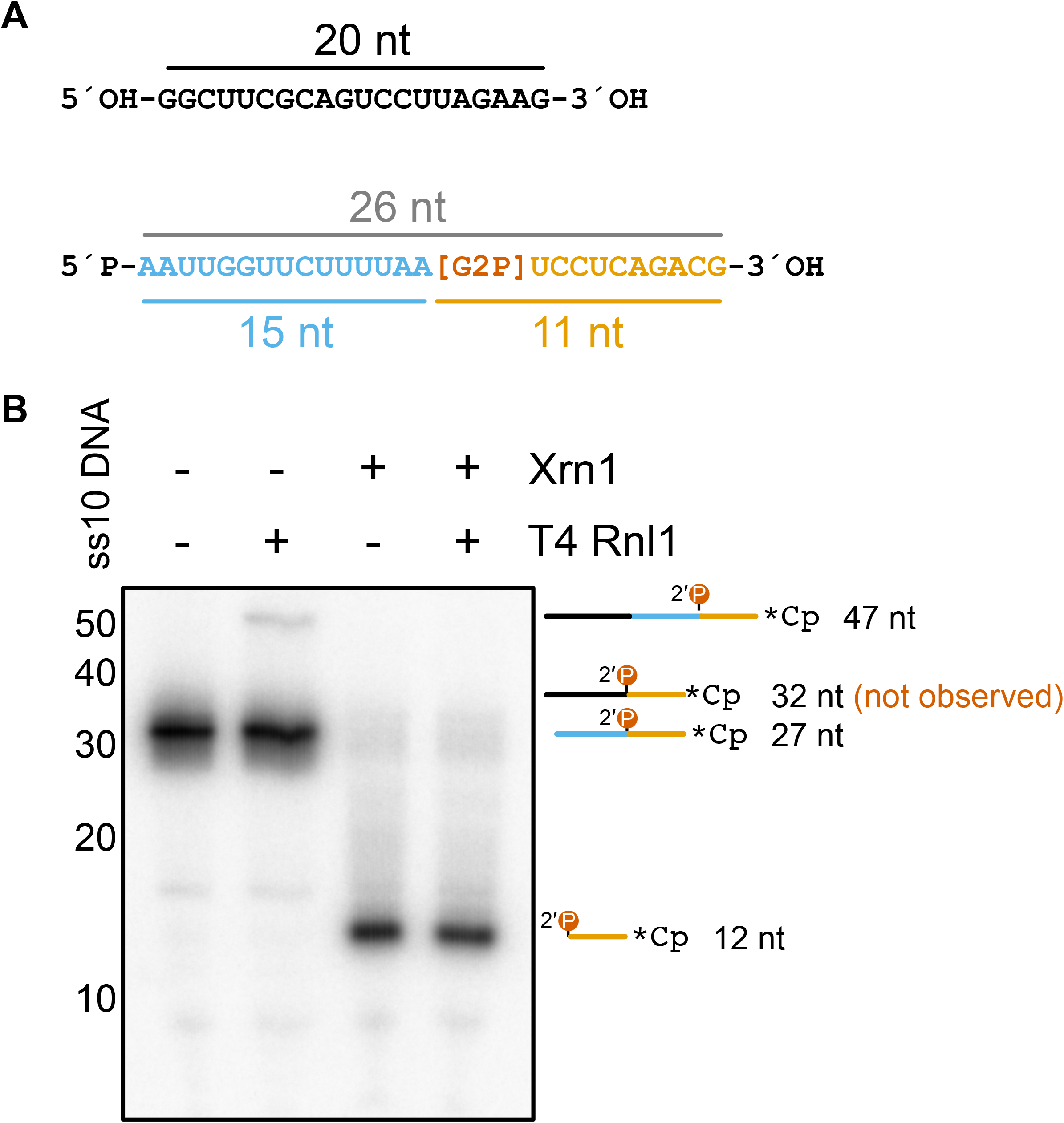
Terminal 5′, 2′-phosphates are resistant to T4 RNL1 ligation. (**A**) Unmodified 20 nt oligo and 2′-phosphorylated 26 nt oligo used in this experiment. (**B**) 26 nt oligos were 3’-end labeled with radiolabeled pCp prior to incubation with or without rXrn1. After digestion, all samples were treated with phenol:chloroform and ethanol precipitated, and then mixed with equimolar amounts of the 20 nt oligo, followed by a 1 hour incubation with or without T4 RNL1. Ligation products are visible in lane 2, but undetectable in lane 4.

**SUPPLEMENTAL FIGURE 2.**
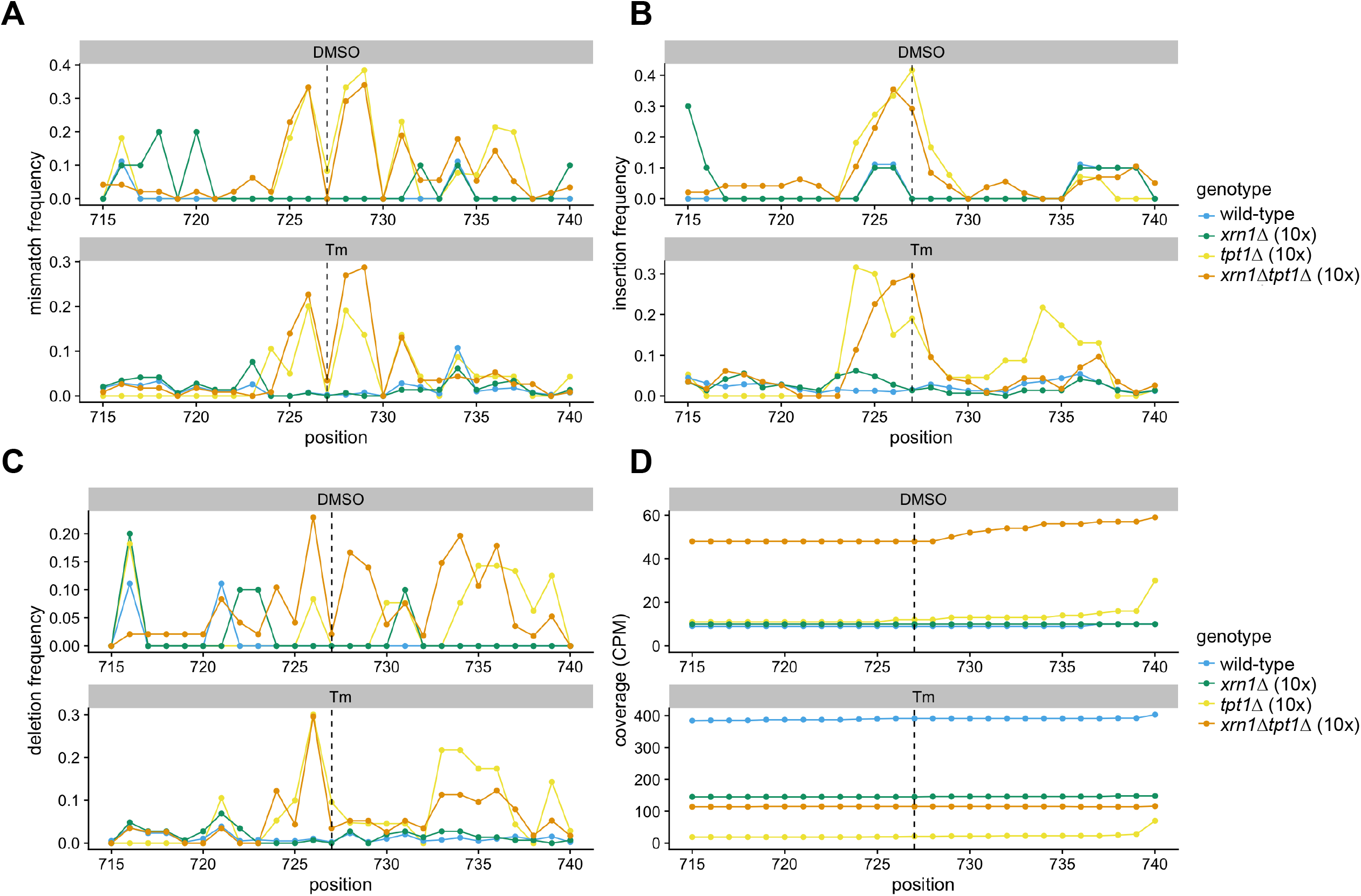
Per-nucleotide base calling “errors” at the *HAC1*^s^ splice junction. (**A**) Mismatch frequency, (**B**) insertion frequency, (**C**) deletion frequency, and (**D**) read coverage in counts per million for eight direct RNA sequencing libraries. In each plot, the 2′-phosphorylated position is indicated by a dashed line.

**SUPPLEMENTAL FIGURE 3.**
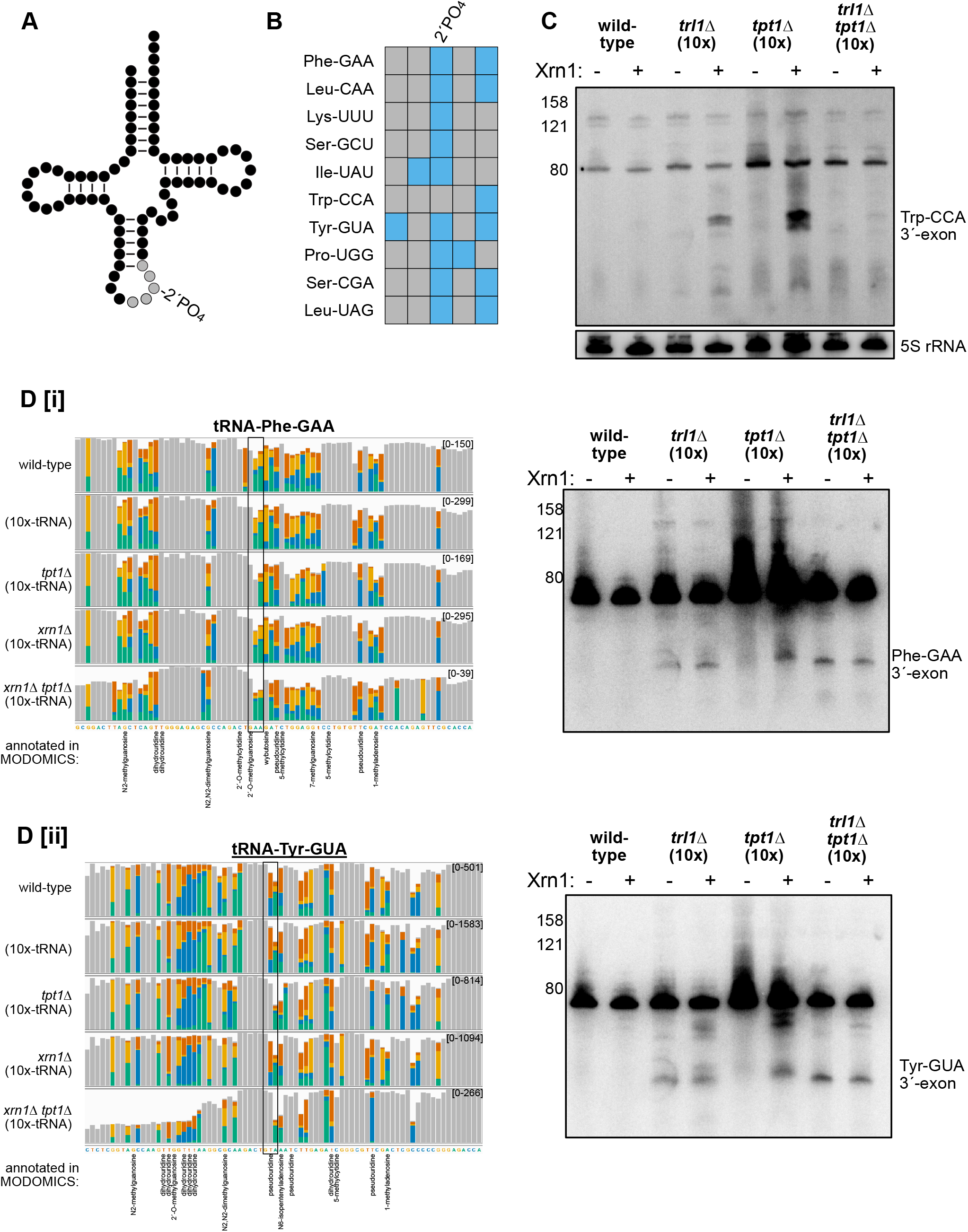
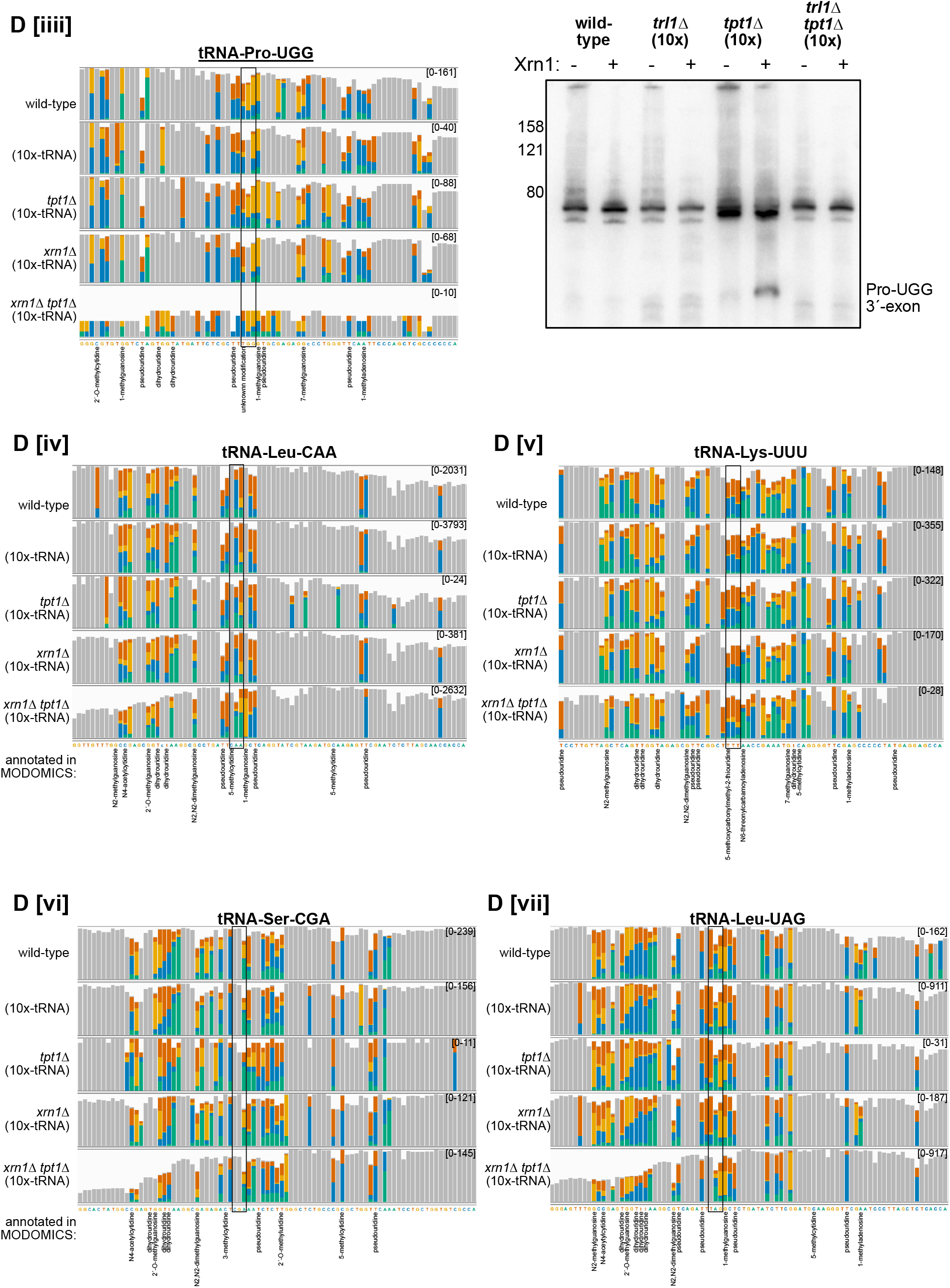
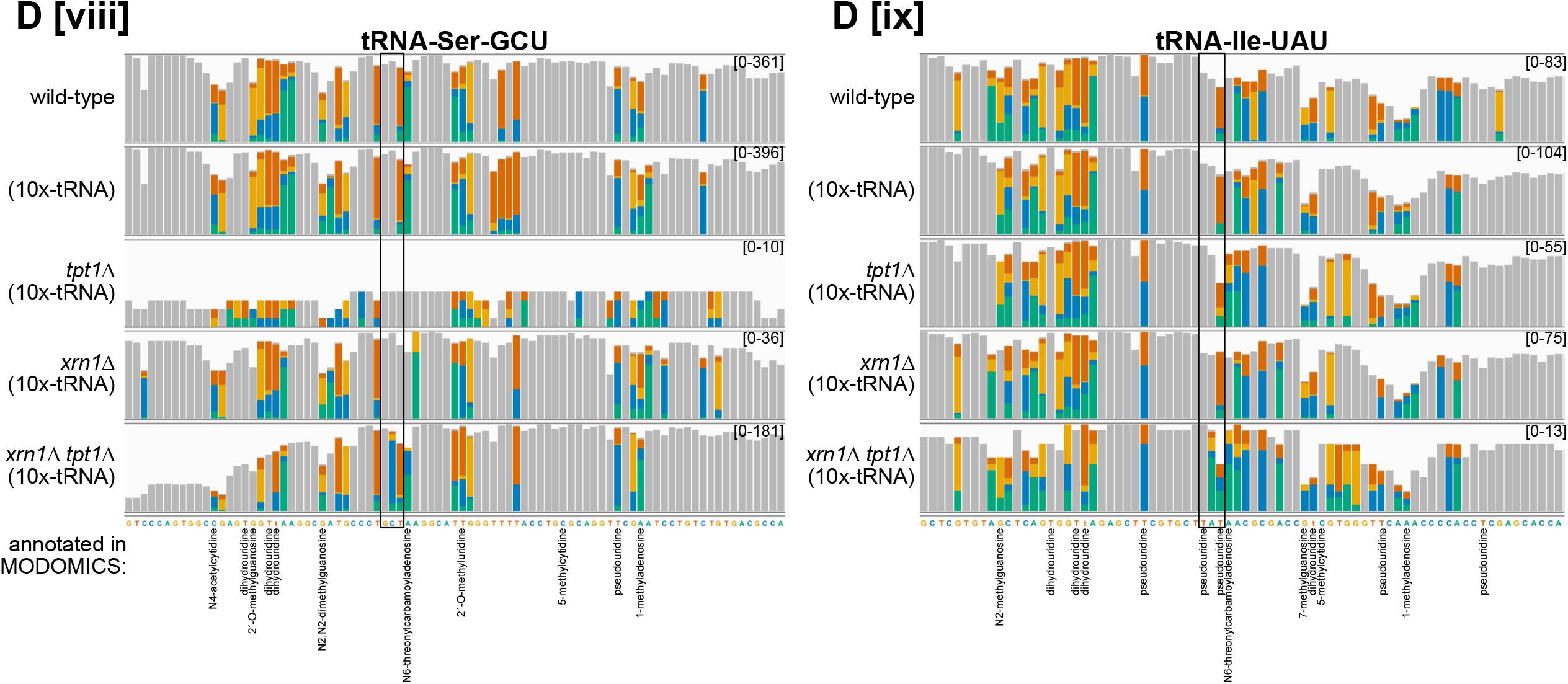
Detection of 2′-phosphates on intron-containing tRNAs is complicated by the presence of additional tRNA modifications. (**A**) Cloverleaf diagram of a spliced tRNA from *tpt1Δ* cells, indicating the 2′-phosphorylated nucleotide located immediately 3′ of the anticodon, and the 5mer centered on this position in gray. (**B**) The RNA modification landscape within the indicated 5mer in (**A**) for each intron-containing tRNA in *S. cerevisiae*. Blue boxes indicate positions with known tRNA modifications in the MODOMICS database. (**C**) The 2′-phosphate on tRNA-Trp-CCA can be detected by northern blotting for 3′-exon in total RNA from RNA repair strains treated with and without recombinant Xrn1 *in vitro*. 3′-exon degradation intermediates accumulate in *tpt1Δ* (10x-tRNA) cells, while two larger exonuclease-resistant bands are also present, albeit less abundant, in RNA from *trl1Δ* (10x-tRNA) cells treated with rXrn1. Given that the MODOMICS database lists annotated methyl groups on this mature tRNA at nucleotides 31 and 33 ^25^, and that 2′-O-methyl groups also inhibit Xrn1^38^, albeit substantially less robustly than 2′-phosphates (**Fig. 1b**), it is possible that these larger bands are produced by 2′-O-methyl-mediated rXrn1 inhibition. However, in the context of a *trl1Δ tpt1Δ* double mutant, these degradation intermediates disappear, raising the possibility of additional crosstalk between tRNA splicing and tRNA modification pathways. In the lower sub-panel, the same membrane has been stripped and reprobed for 5S rRNA as a loading control. (**D**) IGV snapshots of direct tRNA sequencing reads mapping to intron-containing *S. cerevisiae* tRNAs. Each colored bar represents a position with >20% mismatching to the reference base; gray bars indicate positions which did not exceed this threshold. Numbers in brackets at right indicate the Y axis range in read counts, and tRNA anticodons are surrounded by a black box. Modified positions reported in MODOMICS are annotated below the reference sequence. To the right of select IGV alignment views are northern blots of total RNA from *S. cerevisiae* strains treated with and without recombinant Xrn1 *in vitro*, using a 3′-exon hybridizing probe.

**SUPPLEMENTAL FIGURE 4.**
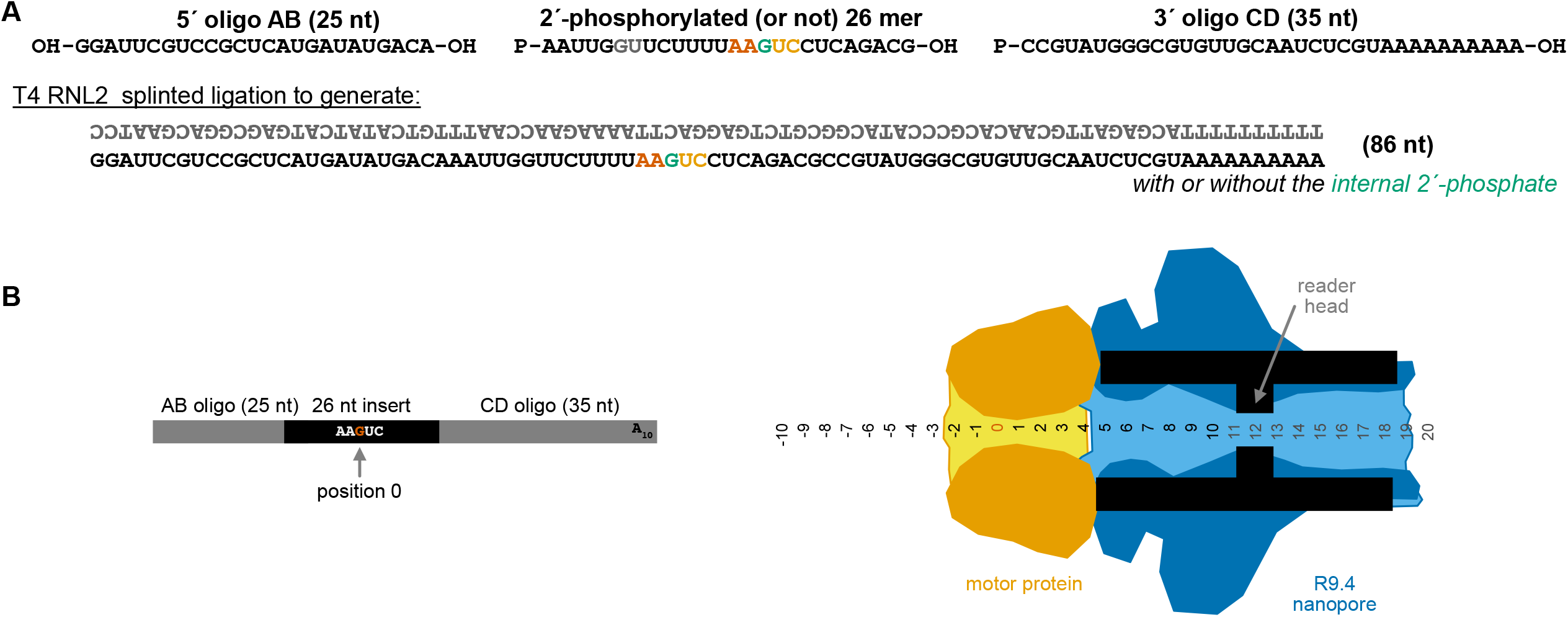
(**A**) Strategy for generation of a 2′-phosphate containing synthetic RNA oligonucleotide for direct RNA sequencing. Flanking oligos AB and CD were ligated to the 2′-phosphorylated or unmodified central 26 nucleotide oligo via a T4 RNL2 splinted ligation, treated with DNase I to destroy the splint, and gel purified to retain only the full 86 nt product, which was prepared for direct RNA sequencing on nanopore beginning with the RTA adapter ligation step. (**B**) Representation of this synthetic oligonucleotide with the 2′-phosphorylated nucleotide as position 0, within the context of an R9.4 nanopore with a docked motor protein. As the modified nucleotide at position 0 is entering the helicase, the nanopore is sensing positions 11-13 nucleotides downstream within the reader head.

**SUPPLEMENTAL FIGURE 5.** Scatter plot of the difference in mean current intensity vs. difference in mean dwell time for modified and unmodified RNAs across a window extending from 10 nt 5′ of the modified position to 20 nt 3′ of the modified site. Each dot represents an individual position across the window of interest.

## Supplemental Tables

**SUPPLEMENTAL TABLE 1.**
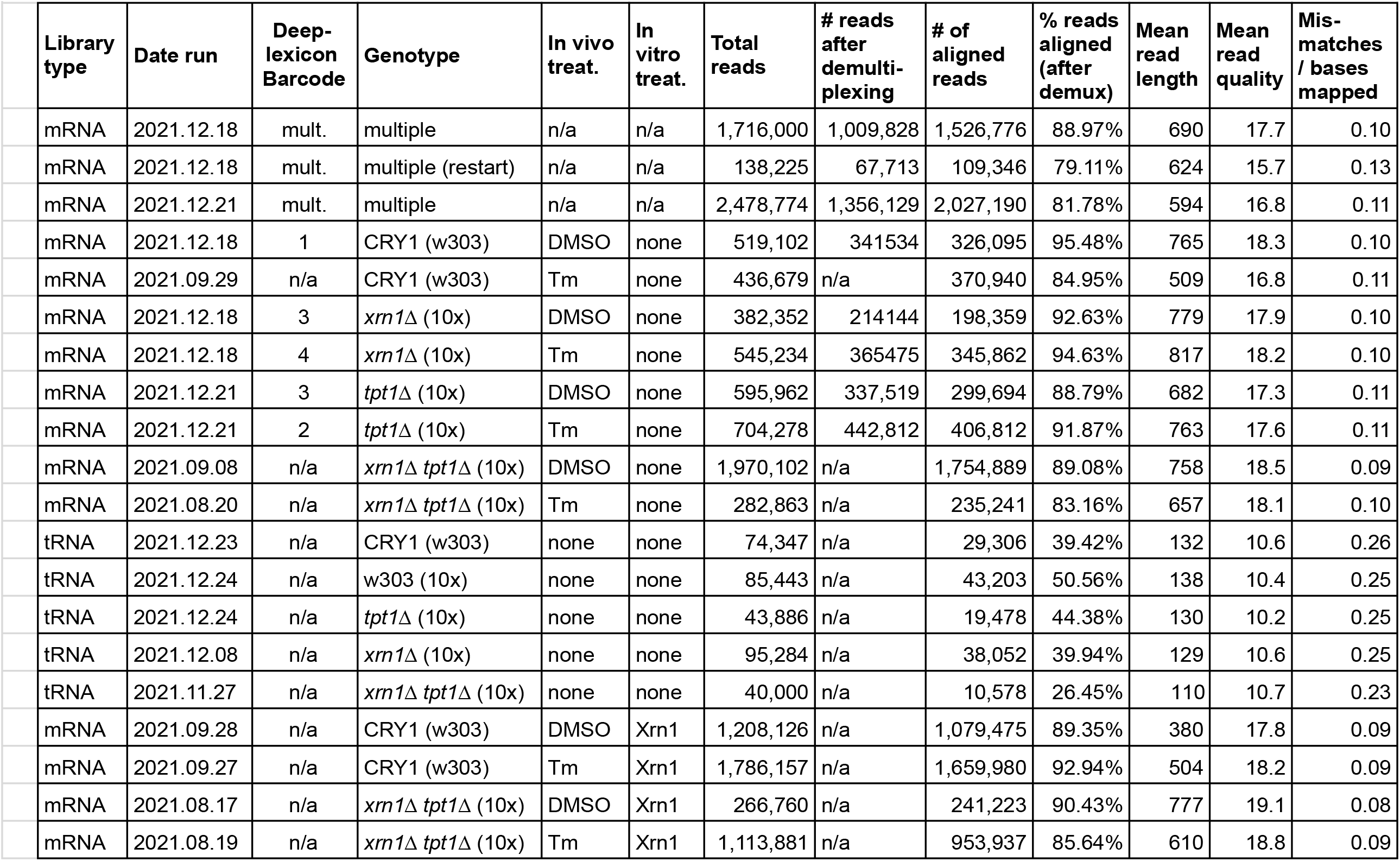
Library metrics and metadata.

| SUPPLEMENTAL TABLE 1 |  |  |  |  |  |  |  |  |  |  |  |  |  |
| --- | --- | --- | --- | --- | --- | --- | --- | --- | --- | --- | --- | --- | --- |
|  | Library type | Date run | Deep-lexicon Barcode | Genotype | In vivo treat. | In vitro treat. | Total reads | # reads after demultiplexing | # of aligned reads | % reads aligned (after demux) | Mean read length | Mean read quality | Mis-matches / bases mapped |
|  | mRNA | 2021.12.18 | mult. | multiple | n/a | n/a | 1,716,000 | 1,009,828 | 1,526,776 | 88.97% | 690 | 17.7 | 0.10 |
|  | mRNA | 2021.12.18 | mult. | multiple (restart) | n/a | n/a | 138,225 | 67,713 | 109,346 | 79.11% | 624 | 15.7 | 0.13 |
|  | mRNA | 2021.12.21 | mult. | multiple | n/a | n/a | 2,478,774 | 1,356,129 | 2,027,190 | 81.78% | 594 | 16.8 | 0.11 |
|  | mRNA | 2021.12.18 | 1 | CRY1 (w303) | DMSO | none | 519,102 | 341534 | 326,095 | 95.48% | 765 | 18.3 | 0.10 |
|  | mRNA | 2021.09.29 | n/a | CRY1 (w303) | Tm | none | 436,679 | n/a | 370,940 | 84.95% | 509 | 16.8 | 0.11 |
|  | mRNA | 2021.12.18 | 3 | *xrn1*Δ (10x) | DMSO | none | 382,352 | 214144 | 198,359 | 92.63% | 779 | 17.9 | 0.10 |
|  | mRNA | 2021.12.18 | 4 | *xrn1*Δ (10x) | Tm | none | 545,234 | 365475 | 345,862 | 94.63% | 817 | 18.2 | 0.10 |
|  | mRNA | 2021.12.21 | 3 | *tpt1*Δ (10x) | DMSO | none | 595,962 | 337,519 | 299,694 | 88.79% | 682 | 17.3 | 0.11 |
|  | mRNA | 2021.12.21 | 2 | *tpt1*Δ (10x) | Tm | none | 704,278 | 442,812 | 406,812 | 91.87% | 763 | 17.6 | 0.11 |
|  | mRNA | 2021.09.08 | n/a | *xrn1*Δ *tpt1*Δ (10x) | DMSO | none | 1,970,102 | n/a | 1,754,889 | 89.08% | 758 | 18.5 | 0.09 |
|  | mRNA | 2021.08.20 | n/a | *xrn1*Δ *tpt1*Δ (10x) | Tm | none | 282,863 | n/a | 235,241 | 83.16% | 657 | 18.1 | 0.10 |
|  | tRNA | 2021.12.23 | n/a | CRY1 (w303) | none | none | 74,347 | n/a | 29,306 | 39.42% | 132 | 10.6 | 0.26 |
|  | tRNA | 2021.12.24 | n/a | w303 (10x) | none | none | 85,443 | n/a | 43,203 | 50.56% | 138 | 10.4 | 0.25 |
|  | tRNA | 2021.12.24 | n/a | *tpt1*Δ (10x) | none | none | 43,886 | n/a | 19,478 | 44.38% | 130 | 10.2 | 0.25 |
|  | tRNA | 2021.12.08 | n/a | *xrn1*Δ (10x) | none | none | 95,284 | n/a | 38,052 | 39.94% | 129 | 10.6 | 0.25 |
|  | tRNA | 2021.11.27 | n/a | *xrn1*Δ *tpt1*Δ (10x) | none | none | 40,000 | n/a | 10,578 | 26.45% | 110 | 10.7 | 0.23 |
|  | mRNA | 2021.09.28 | n/a | CRY1 (w303) | DMSO | Xrn1 | 1,208,126 | n/a | 1,079,475 | 89.35% | 380 | 17.8 | 0.09 |
|  | mRNA | 2021.09.27 | n/a | CRY1 (w303) | Tm | Xrn1 | 1,786,157 | n/a | 1,659,980 | 92.94% | 504 | 18.2 | 0.09 |
|  | mRNA | 2021.08.17 | n/a | *xrn1*Δ *tpt1*Δ (10x) | DMSO | Xrn1 | 266,760 | n/a | 241,223 | 90.43% | 777 | 19.1 | 0.08 |
|  | mRNA | 2021.08.19 | n/a | *xrn1*Δ *tpt1*Δ (10x) | Tm | Xrn1 | 1,113,881 | n/a | 953,937 | 85.64% | 610 | 18.8 | 0.09 |

**SUPPLEMENTAL TABLE 2.**
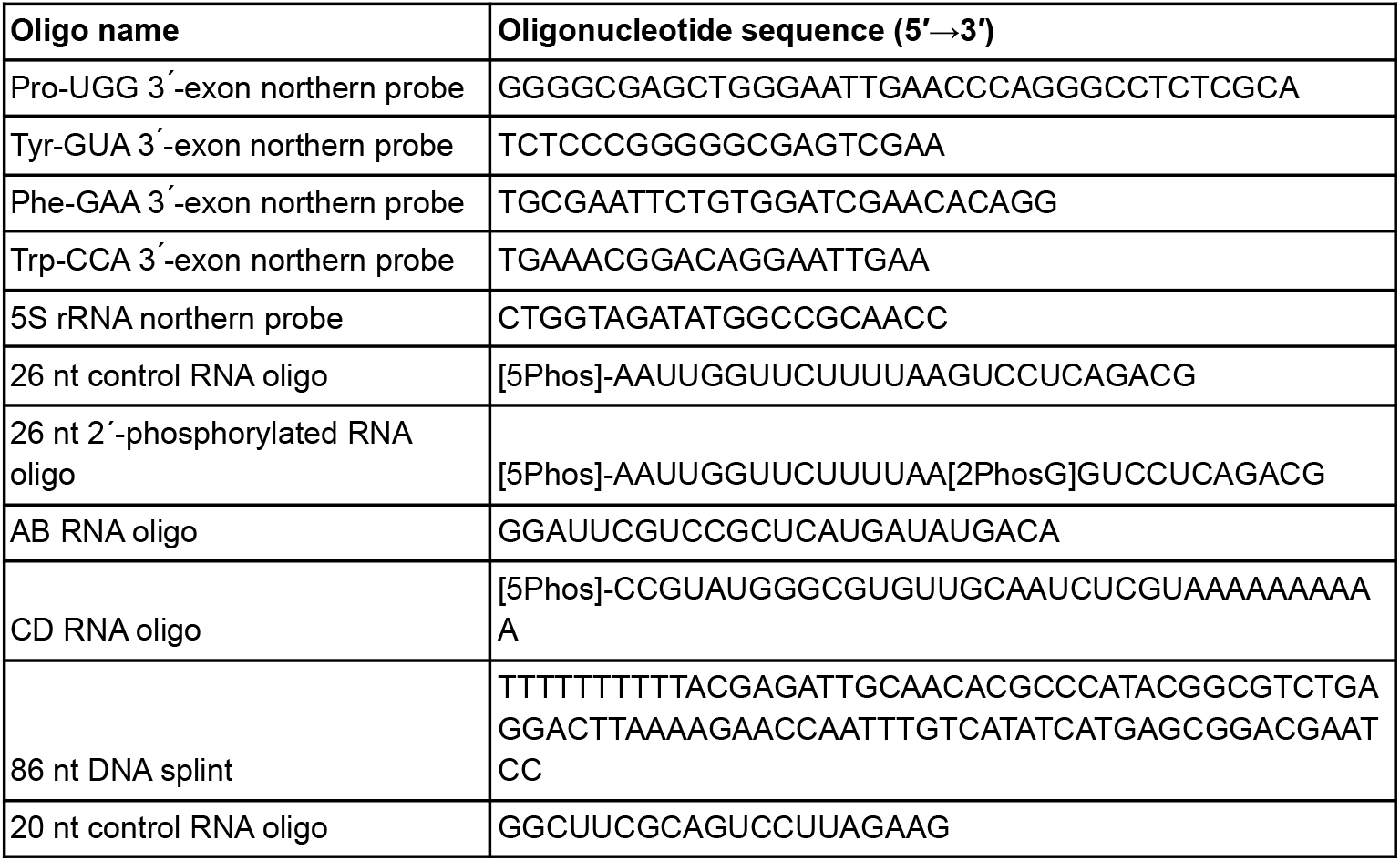
Oligonucleotides used in this study.

**SUPPLEMENTAL TABLE 3.** Yeast strain numbers and genotypes.

| Strain ID | Genotype | Source |
| --- | --- | --- |
| YJH682 | wild-type (CRY1) | Mingxia Huang |
| YJH932 | (pAG424-10x-tRNA) |  |
| YJH632 | *xrn1Δ::HygMX* |  |
| YJH834 | *tpt1Δ::LEU2* (pAG424-10x-tRNA) |  |
| YJH902 | *tpt1Δ::LEU2 xrn1Δ::HygMX* (pAG424-10x-tRNA) |  |
| YJH835 | *trl1Δ::KanMX* (pAG424-10x-tRNA) |  |
| YJH927 | *trl1Δ::KanMX tpt1Δ::LEU2* (pAG424-10x-tRNA) |  |

## Data availability

Raw data will be deposited at NCBI GEO and a pipeline for analysis is available at https://github.com/hesselberthlab/RNARePore.

## Acknowledgements

We thank N. Mukherjee for comments on the manuscript. This work was supported by the National Institutes of Health (R35 GM119550 and T32 GM136444), the Molecular Biology Program and Bolie Family Foundation (L.W.) and the RNA Bioscience Initiative (L.W.)

